# Machine learning-based approach KEVOLVE efficiently identifies SARS-CoV-2 variant-specific genomic signatures

**DOI:** 10.1101/2022.02.07.479343

**Authors:** Dylan Lebatteux, Hugo Soudeyns, Isabelle Boucoiran, Soren Gantt, Abdoulaye Baniré Diallo

## Abstract

Machine learning was shown to be effective at identifying distinctive genomic signatures among viral sequences. These signatures are defined as pervasive motifs in the viral genome that allow discrimination between species or variants. In the context of SARS-CoV-2, the identification of these signatures can assist in taxonomic and phylogenetic studies, improve in the recognition and definition of emerging variants, and aid in the characterization of functional properties of polymorphic gene products. In this paper, we assess KEVOLVE, an approach based on a genetic algorithm with a machine-learning kernel, to identify multiple genomic signatures based on minimal sets of *k*-mers. In a comparative study, in which we analyzed large SARS-CoV-2 genome dataset, KEVOLVE was more effective at identifying variant-discriminative signatures than several gold-standard statistical tools. Subsequently, these signatures were characterized using a new extension of KEVOLVE (KANALYZER) to highlight variations of the discriminative signatures among different classes of variants, their genomic location, and the mutations involved. The majority of identified signatures were associated with known mutations among the different variants, in terms of functional and pathological impact based on available literature. Here we showed that KEVOLVE is a robust machine learning approach to identify discriminative signatures among SARS-CoV-2 variants, which are frequently also biologically relevant, while bypassing multiple sequence alignments. The source code of the method and additional resources are available at: https://github.com/bioinfoUQAM/KEVOLVE.

## Introduction

Severe acute respiratory syndrome coronavirus (SARS-CoV-2) is the etiological agent of coronavirus disease 2019 (COVID-19). This highly infectious coronavirus was first identified in December 2019 in Wuhan, China [1]. It belongs to the betacoronavirus genus, which includes SARS-CoV-1 and Middle East respiratory syndrome-related coronavirus (MERS-CoV) [2]. The genome of SARS-CoV-2 is a single-stranded RNA molecule composed of approximately 30,000 nucleotides. The nucleotide sequence identity of SARS-CoV-2 with SARS-CoV-1 and MERS-CoV is 79.5% and 50%, respectively [3, 4]. The SARS-CoV-2 genome encodes 29 different proteins, including 16 nonstructural proteins, 4 structural proteins, and 9 accessory proteins (see Fig 1 adapted from [5]). The N (nucleocapsid) protein contains the viral RNA genome, while the S (spike), E (envelope), and M (membrane) proteins together form the viral envelope [6]. SARS-CoV-2 exhibits a notably high mutation rate, with numerous mutations—particularly in the spike gene—correlated to increased SARS-CoV-2 transmission rates [7], augmented fusogenic and pathogenic properties of the virus [8], as well as the emergence of new variants that could diminish the efficacy of existing COVID-19 vaccines and antibody-based therapies [9].

**Fig 1.**
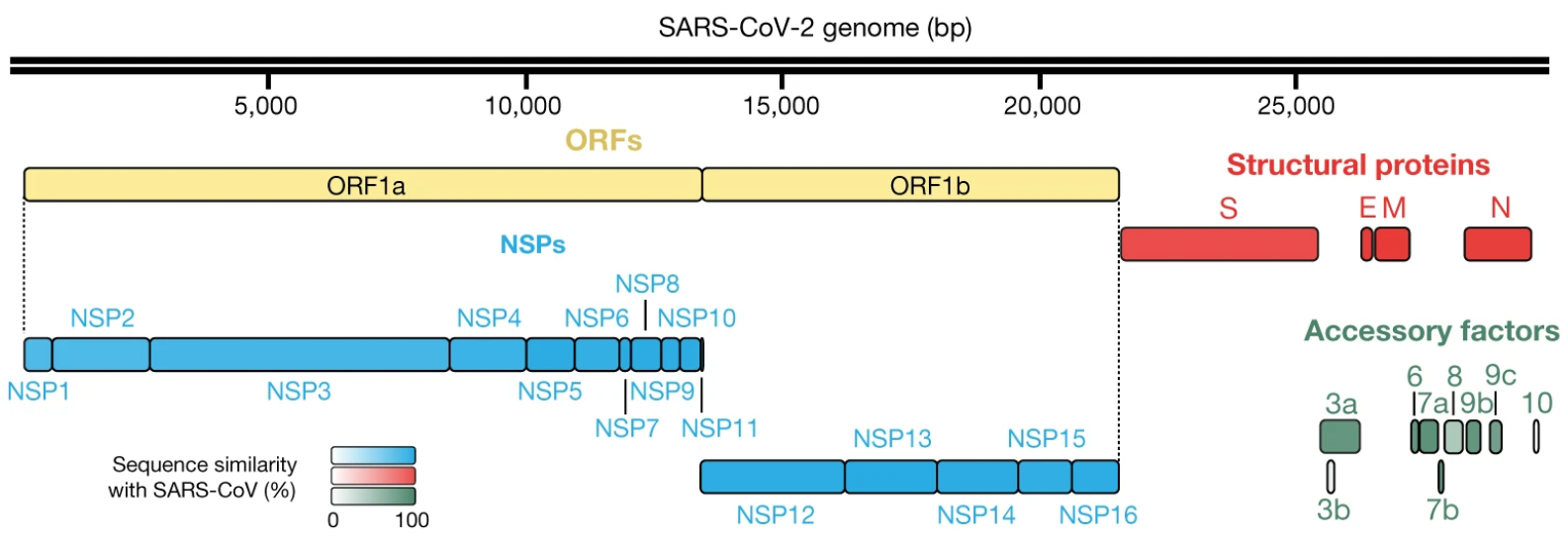
SARS-CoV-2 genome organization. Four structural proteins (red), 16 non-structural proteins (NSPs; blue), and 9 accessory factors (green) are shown. ORFs (open reading frames; yellow) 1a and 1b encode polyproteins. The protein sequence similarity with SARS-CoV homologues (when homologues exist) is depicted by the color intensity.

Given its rapid rate of evolution, it is important to be able to efficiently identify genomic signatures that can distinguish between different variants of SARS-CoV-2 and highlight potential functional changes. These signatures, also known as species- or variant-specific motifs that are prevalent throughout the viral genome [10], can contribute to taxonomic [11] and phylogenetic [12] studies to differentiate distinct groups of variants, provide insight into their evolutionary history [10], help to understand the structure of the viral quasispecies [13], and facilitate mechanistic studies to determine the functional basis of variant-specific differences in virulence [14]. To identify discriminative motifs, or genomic signatures, among different groups of biological sequences, the traditional approach is to compute multiple sequence alignments using tools such as MUSCLE [15], Clustal W/X [16], or MAFFT [17]. These alignments are then analyzed to identify divergent genomic regions that constitute the discriminative motifs. However, multiple alignment approaches have significant limitations when applied to viral genomes [14].

First, alignment-based approaches are generally computationally and time-intensive, making them less well suited for dealing with large viral sequence datasets that are increasingly available [18]. In fact, computing an accurate multi-sequence alignment is an NP-hard problem with (2*N*)!*/*(*N* !)^2^ possible alignments for two sequences of length *N* [19], which means that in some cases, the alignment cannot be solved within a realistic time frame or involves significant compromise in accuracy [17]. Even with dynamic programming, the time requirement is on the order of the product of the lengths of the input sequences [20]. Second, alignment algorithms assume that homologous sequences consist of a series of more or less conserved linearly arranged sequence segments. However, this assumption, named collinearity, is often questionable, especially for RNA viruses [21]. This is because RNA viruses show extensive genetic variation due to high mutation rates, as well as high frequencies of genetic recombination, horizontal gene transfer, and gene duplication, leading to the gain or the loss of genetic material [22]. Finally, performing multiple alignments often requires adjusting several parameters, such as substitution matrices, deviation penalties, and thresholds for statistical parameters, which are dependent on prior knowledge about the evolution of the compared sequences [21]. However, the adjustment of these parameters is sometimes arbitrary and requires a trial-and-error approach, and research has shown that small variations in these parameters can significantly impact the quality of alignments [23].

To address the limitations of discriminative motif identification using multiple sequence alignment, specialized statistical-based tools were developed, such as MEME [24, 25]. MEME has a discriminative mode [26] that identifies enriched motifs that distinguish a primary set of sequences from a control set. Other MEME tools were also developed, including STREME [27], the most powerful tool for discovering motifs in sequence datasets. STREME uses a generalized suffix tree and evaluates motifs using a statistical test that compares the enrichment of matches to the motif in the primary set of sequences to the control set [27].In recent years, a series of machine-learning techniques were developed and widely used in the field of genomics, and were proven to be highly effective for solving complex and large-scale data analysis problems [28]. For example, the CASTOR study [29] demonstrated the usefulness of machine learning models coupled with restriction fragment length polymorphism (RFLP) signatures for classifying viral genomic sequences, achieving f1-scores *≥* 0.99 for predicting hepatitis B virus and human papillomavirus genomes. However, these signatures were found to have limitations in predicting HIV sequences, resulting in an f1-score *≤* 0.90. To address this issue, the KAMERIS study [30] used *k*-mers (nucleotide subsequences of length *k*) to characterize the sequences provided to the learning model. To reduce the exponential number of features (4^*k*^) associated with *k*-mers, KAMERIS applied truncated singular value decomposition for dimensionality reduction, but this transformation affected the ability to identify and analyze relevant features identified by the machine-learning model for discriminating between groups of sequences.

In response to this challenge, CASTOR-KRFE [31] was developed as a method for identifying minimal sets of genomic signatures based on minimal sets of *k*-mers to discriminate among multiple groups of genomic sequences. During cross-validation evaluations covering a wide range of viruses, CASTOR-KRFE successfully identified minimal sets of motifs, which when combined with supervised learning algorithms, resulted in average f1-scores *≥* 0.96 [31]. However, this study was limited to identifying the optimal set of motifs, rather than exploring suboptimal sets in the feature space, which can be a major limitation when dealing with viral sequences with high genomic diversity or when attempting to infer biological functions based on the identified motifs. To overcome this limitation, KEVOLVE [32] was developed as a new method that uses a genetic algorithm incorporating a machine-learning kernel to identify multiple minimal subsets of discriminative motifs. A preliminary comparative study on HIV nucleotide sequences showed that the KEVOLVE-identified motifs allowed for the construction of models that outperformed specialized HIV prediction tools [32]. In the context of the COVID-19 pandemic, this paper assessed the performance of KEVOLVE in a comparative study with several reference tools (MEME, STREME, and CASTOR-KRFE) for identifying discriminative motifs in the genomes of SARS-CoV-2 variants. The identified motifs were then analyzed using the new KEVOLVE extension (KANALYZER) to extract the associated information, and this information, which is discussed in light of the available literature to highlight the potential biological functions of the sequences/motifs in question.

## Materials and methods

To assess the accuracy of KEVOLVE in identifying discriminative motifs, we conducted a comparative study with specialized tools. This involved using each tool to identify a subset of discriminating motifs in a set of training sequences of SARS-CoV-2 variants. These sets of motifs were designed to provide genomic signatures specific to each variant. In a second step, we used these signatures and a supervised learning algorithm to fit a prediction model on the training sequences. Then, we evaluated the quality of the signatures by predicting the trained models on a large test set of unknown sequences. Finally, we used KANALYZER, the latest extension of KEVOLVE, to analyze the variant-discriminative motifs identified by KEVOLVE and assess their potential functional impact based on their location in the genome, as previously described in the literature.

### Discriminative motif identification tools

We first evaluated KEVOLVE [32], a machine learning method based on a genetic algorithm for identifying multiple minimal sets of *k*-mers to discriminate nucleotide sequences. KEVOLVE takes as input a set of labeled nucleotide sequences and a parameter *k*, which corresponds to the length of the *k*-mers used to represent the sequences in an occurrence matrix. KEVOLVE starts by using a meta-transformer to remove *k*-mers with low discriminative contribution based on importance weights assigned by a linear Support Vector Machine (SVM). Then, the genetic algorithm begins its search by initializing several subsets (chromosomes) composed of a reduced set of *k*-mers (genes). Each chromosome is evaluated in a cross-validation process where prediction models are trained and tested on nucleotide sequences represented by the genes in the chromosome. The chromosomes with the best scores are then subjected to mutation/crossover processes. The mutation process involves randomly substituting a gene with another within a chromosome, and the crossover process involves exchanging genes between different chromosomes. In addition, the genes in the best chromosome have an increased probability of being selected in the next iteration. The next generation is then composed of the best current chromosomes and new chromosomes, which are generated based on the updated probability of selection. This process is repeated and coupled with a progressive increase in chromosome size until a stopping criterion is met (number of iterations or performance score of the solutions). The detailed KEVOLVE pseudo code is available in the original article [32], and the algorithm code can be accessed in the GitHub repository.

The second tool we evaluated was CASTOR-KRFE [31], an alignment-free machine learning approach for identifying a set of genomic signatures based on *k*-mers to discriminate between groups of nucleic acid sequences. The core of CASTOR-KRFE is based on feature elimination using SVM (SVM-RFE). It identifies the optimal length of *k* to maximize classification performance and minimize the number of features, providing a solution to the problem of identifying the optimal length of *k*-mers for genomic sequence classification [33]. The third tool we evaluated was MEME (discriminative mode) [26], a tool from the MEME suite [25] specialized in motif identification. MEME takes two sets of sequences as input and identifies enriched motifs that discriminate the primary set from the control set. By default, MEME assumes that all positions in the sequences have an equal chance of being a motif site. However, in discriminative mode, the algorithm uses additional information such as sequence conservation, nucleosome positioning, and negative examples to compute a measure of the probability that a discriminative motif starts at each position in each sequence [26].This measure, called “position specific prior” (PSP), is then used to guide the sequence motif discovery algorithm in the primary set, resulting in motifs that are more likely to discriminate it from the control set [34]. MEME also allows for the specification of a potential motif distribution type to improve the sensitivity and quality of the motif search. There are two available options in discriminative mode: zero or one occurrence per sequence (ZOOPS), where MEME assumes that each sequence may contain at most one occurrence of each motif, and one occurrence per sequence (OOPS), where MEME assumes that each sequence in the dataset contains exactly one occurrence of each motif. The last tool we evaluated was STREME [27], which was found to be more accurate, sensitive, and thorough than several widely used algorithms in a recent comparative study [27]. STREME’s algorithm uses a data structure called a generalized suffix tree and evaluates motifs using a one-sided statistical test of the enrichment of matches to the motif in a primary set of sequences compared to a control set. STREME assumes that each primary sequence may contain ZOOPS of the motif, but the discovery of the motif will not be negatively affected if a primary sequence contains more than one occurrence.

### Dataset

To set up the most comprehensive evaluation framework possible, we built a dataset of 334,956 SARS-CoV-2 genomes representing the different variants defined by the World Health Organization (WHO) with at least 100 available sequences. The sequences for this dataset, covering variants Alpha (B.1.1.7), Beta (B.1.351), Gamma (P.1), Delta (B.1.617.2), Kappa (B.1.617.1), Epsilon (B.1.427/B.1.427), Iota (B.1.526), Eta (B.1.525), Lambda (C.37), and Omicron (B.1.1.529/BA.x), were downloaded on November 1, 2022 from the NCBI database [35] using their command line data download tool (https://www.ncbi.nlm.nih.gov/datasets/docs/v2/how-tos/virus/get-sars2-genomes/). We only included complete genomes with high coverage (less than 1% missing nucleotides) in our dataset (Table 1), and the list of accession ids for the sequences used in our different datasets is available on our GitHub repository.

**Table 1.** Genomic sequence dataset of SARS-CoV-2 variants.

| WHO Label | Pango Lineage | Country of origin | Number of sequences |
| --- | --- | --- | --- |
| Alpha | B.1.1.7 | United Kingdom | 175,212 |
| Beta | B.1.351 | South Africa | 695 |
| Gamma | P.1 | Brazil | 8,129 |
| Delta | B.1.617.2 | India | 9,408 |
| Kappa | B.1.617.1 | India | 127 |
| Epsilon | B.1.427/B.1.429 | USA (California) | 14,674 |
| Iota | B.1.526 | USA (New York) | 19,274 |
| Eta | B.1.525 | United Kingdom/Nigeria | 716 |
| Lambda | C.37 | Peru | 428 |
| Omicron | B.1.1.529/BA.x | South Africa (Botswana) | 106,293 |
| **Total number of sequences** |  |  | **334,956** |

### Benchmarking

We assessed the performance of the different tools to identify discriminative motifs using an established approach [31]. We performed a repeated *K*-fold evaluation 100 times with a different randomization at each repetition. For each iteration, 2,500 sequences were used to form a training set and the rest (332,456) were used as a testing set. In the training set, the variants were represented by 250 sequences, with the exception of Kappa, which was represented by 100 sequences due to the low number of available sequences. Alpha and Omicron were each represented by 350 and 300 sequences, respectively, due to the large number of available sequences. At each iteration, the training sets were given as input to each tool to identify the motifs that discriminate the sequences of the variants. The identified motifs, along with the training sequences, were used to train a machine-learning algorithm (linear-SVM). This model was then used to predict the test set, and different performance metrics were calculated. For each iteration, we computed the unweighted average of precision, recall, and f1-score. By computing each metric as an unweighted average, we avoided the dominance effect of prevalent variants, as demonstrated in Equations 1, 2, and 3.

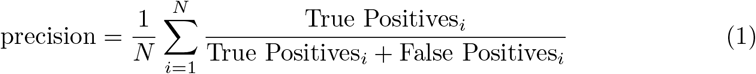

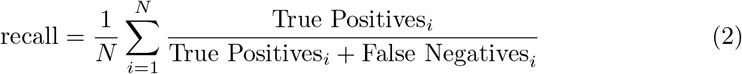

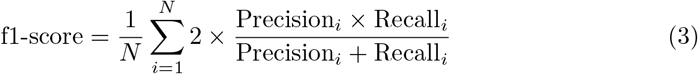

The distributions of the different performance metrics for each tool are illustrated through violin plots in Fig 2.A-C. In addition, to visualize the prediction by class more specifically, we computed the average confusion matrix with its standard deviation for each tool (Fig 2.E-I). Finally, Fig 2.D illustrates the average number of unique motifs identified by each tool during the hundred iterations to train the prediction models.

**Fig 2.**
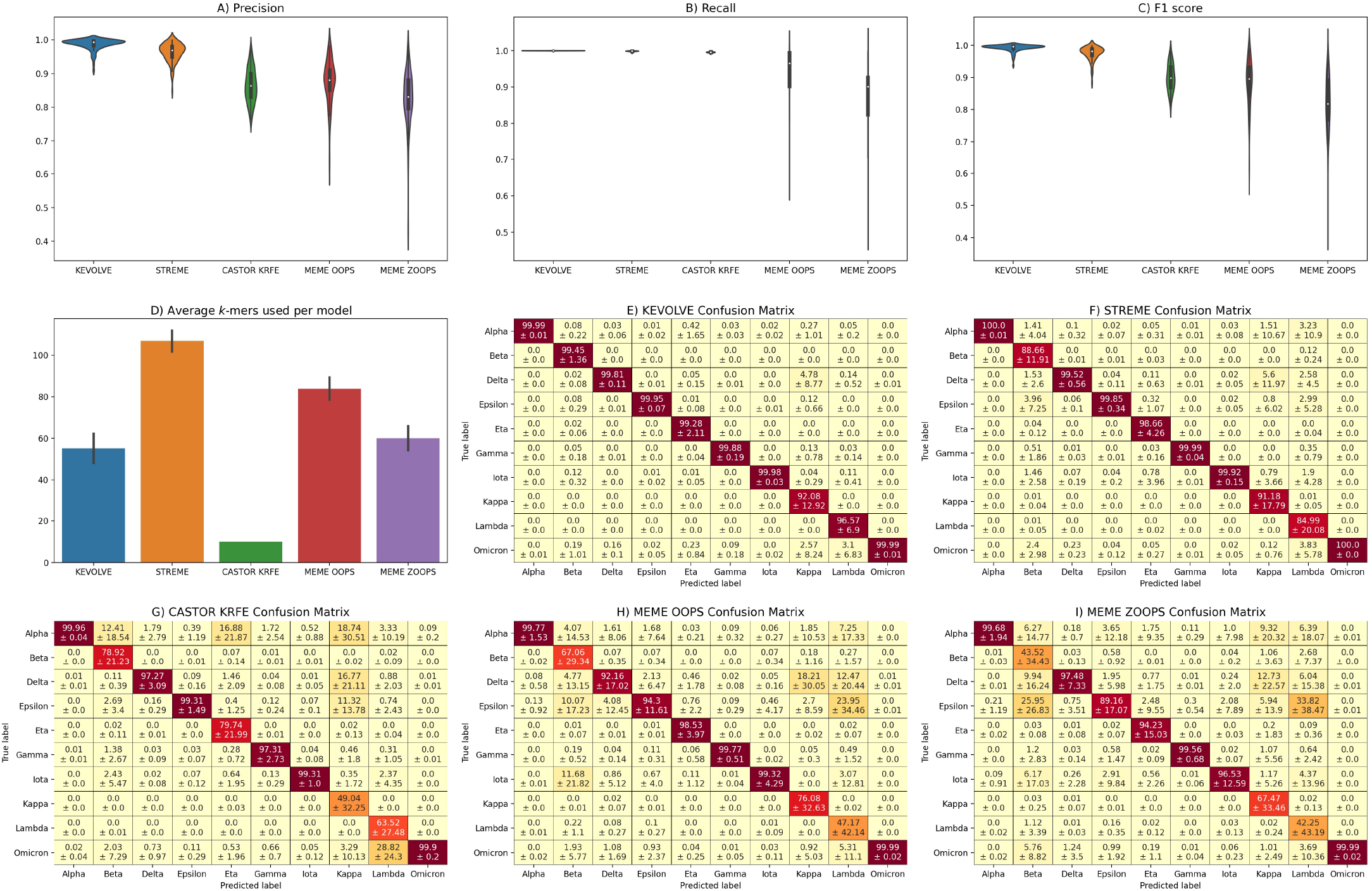
Results of the comparative study. A-C) The violin plots illustrate the distributions of the performance metrics, including Precision, Recall, and F1-score, obtained for the test set predictions during the cross-validation evaluation of 100 iterations. D) The bar plot depicts the average number of motifs identified by each approach to build their prediction model. The black vertical bar indicates the standard deviation. E-I) The confusion matrices represent the average prediction performance as a function of the different variants for each tool over the 100 iterations. Each cell shows the average percentage of the assigned instance in the top value, and the standard deviation in the bottom value.

### Identification of discriminating motifs and tool settings

In the identification phase of the discriminative motifs, we set the length of the motifs to *k* = 9 for two reasons. First, this length is consistent with other studies that have used *k*-mers for viral sequence classification [10, 31, 33]. Second, the selection of a multiple of 3 is consistent with the codon size, and as we use sliding windows with a step of 1 to calculate the number of *k*-mers, encompassing all reading frames, we believe this method facilitates the capture of potential amino acid-level mutations. For KEVOLVE, we set the following search parameters: *n_chromosomes* = 100 (the number of chromosomes generated at each iteration), and *n_genes* = 1 (the number of genes composing the chromosome in the first generation). Initiating with a unitary instance allows KEVOLVE to ascertain the optimal size during its search process since this is unknown, and the training sets vary throughout the evaluation. The stopping criterion parameters were set at *n_iterations* = 1000 and *n solutions* = 10. We utilized the default crossover and mutation rates from a previous study [32] for these parameters. For CASTOR-KRFE, we set the performance threshold to be maintained while reducing the number of features to *T* = 0.99.

To evaluate MEME, considering its limitation to take as input a binary set, we implemented the following process: for each variant *v* in the training set *V*, we selected all sequences belonging to *v* to form the primary set and used the remaining sequences in *V* to form the control set. We then applied MEME to discover motifs that discriminated the primary set from the control set. This process was repeated for each variant *v* in order to build a set of motifs that could discriminate each variant from the others. This set of motifs was used to train a model and predict the testing set in the same configuration as CASTOR-KRFE and KEVOLVE. Both the ZOOPS and OOPS options were evaluated for the associated distribution site parameters. Additionally, to strongly characterize the different groups of sequences, we performed experiments to discover 10 motifs of width 9 for each variant. This choice allows us to theoretically characterize each training set with 100 motifs, assuming there are no duplicates. We applied the same iterative process for identifying motifs to STREME. As mentioned previously, STREME does not require an input parameter for the motif distribution type and handles this automatically. Moreover, considering the number of experiments involved in evaluating the tools of the MEME suite because of their limitation to not handle multi-class sequences, it was not feasible to perform it on their web platform. To handle this, we set up virtual Linux environments where we installed the MEME suite version 5.5.0 with all the necessary dependencies for its functioning. Then several Shell/Python scripts were developed to run the different experiments and process the output files to extract the identified motifs. Finally, we specified that for the tools that identify multiple sets of motifs (KEVOLVE, MEME and STREME), the union of the motifs is used to represent the sequences through the feature matrix at each iteration.

### Analysis of the biological significance of the motifs identified by KEVOLVE

To broaden the utility of KEVOLVE beyond identifying discriminative motifs and building prediction models for nucleotide sequences, we developed KANALYZER [36]. KANALYZER is an extension of KEVOLVE that uses pairwise alignment and parallel computing. It takes as input a reference sequence in GenBank format, a list of nucleotide sequences labeled by their classes related to the organism of the reference sequence, and a list of discriminative motifs associated with the studied sequences. KANALYZER aims to understand the reasons behind a motif’s discriminatory potential by identifying the variations associated with it within different groups of variants. A variation is defined as a nucleotide sequence derived from an initial *k*-mer that has undergone one or more nucleotide changes. KANALYZER generates a report for each motif as output, containing information on their variations that occur in the different nucleotide sequences, their genomic localization, their frequencies of appearance according to the different types of variants, and the resulting mutations at the amino acid level in the case of coding regions. In this study, we used the KANALYZER extension to extract information associated with the discriminative motifs identified by KEVOLVE. The information was derived from the 334,956 sequences we collected and used the SARS-CoV-2 reference sequence NC 045512.2 (Wuhan-Hu-1 isolate, complete genome) for analysis.

## Results and Discussion

### Prediction performances

Initially, we examined the number of discriminative motifs identified by each tool, as summarized in Fig 2.D. CASTOR-KRFE identified the lowest average number of motifs at 10 per iteration, which is minimally constrained by the number of classes in the input dataset. KEVOLVE, MEME ZOOPS, and MEME ZOOPS identified an average of 55, 60, and 84 motifs, respectively. Finally, STREME identified the highest average number of motifs at 107 per iteration, including several degenerate motifs that were converted into classical motifs. The predictive performance of the models based on the motifs identified by each tool is shown in Fig 2.A, 2.B, and 2.C in terms of precision, recall, and f1-score, respectively. KEVOLVE performed the best, with an average score of 0.99 across all metrics. The associated confusion matrix (Fig 2.E) for KEVOLVE indicates that misclassifications sometimes occur, with Kappa sequences being incorrectly predicted as Delta and Omicron in 4.8% and 2.6% of cases, on average. For Lambda variants, approximately 3.1% of the sequences were incorrectly predicted as Omicron. STREME models, which are based on approximately twice as many motifs as KEVOLVE, yielded the second-best predictions with an average performance of 0.96, 1.00, and 0.98 for precision, recall, and f1-score, respectively. The associated confusion matrix for STREME (Fig 2.F) revealed some limitations for Lambda sequences, with more than 12.5% of them being incorrectly predicted as Alpha, Delta, Epsilon, or Omicron, on average. There was also an average of 9% of Beta sequences that were misclassified in a similar manner as Lambda sequences. Like KEVOLVE, STREME models had difficulty predicting certain Kappa variant sequences (≈ 9% on average), with many of them being incorrectly assigned as Delta.

The CASTOR-KRFE method had an average precision of 0.86, a recall of 1.00, and an f1-score of 0.90. The confusion matrix for the CASTOR-KRFE method (Fig 2.G) indicates that it shares the same challenges as the KEVOLVE and STREME methods in inaccurately classifying some Kappa variant sequences, with 19% of these sequences being incorrectly predicted as Alpha, 17% as Delta, and 11% as Epsilon, on average. There were also limitations in the classification of Lambda variants, with nearly 29% of the sequences being incorrectly assigned to Omicron. In addition, more than 12% of the Beta variant sequences were incorrectly assigned to Alpha, on average, and 17% of the Eta variant sequences were incorrectly assigned to Alpha. The MEME OOPS and MEME ZOOPS models showed the poorest prediction performance, with average precisions of 0.97 and 0.83, average recalls of 0.94 and 0.88, and average f1-scores of 0.88 and 0.82, respectively. Both models frequently made classification errors with Lambda variants, which were often incorrectly predicted to be Epsilon. Beta variants were sometimes incorrectly predicted to be Epsilon, Delta, or Iota, and Kappa variants were often incorrectly predicted to be Delta. More detailed results can be seen in the confusion matrices shown in Fig 2.H and 2.I.

### Biological significance of KEVOLVE-identified motifs

To extract biological information related to the motifs identified by KEVOLVE, we first combined all the motifs identified during different iterations. We used these motifs to represent all 334,956 sequences in our dataset and trained a SVM model. We ranked the motifs based on their discriminant contribution, as determined by the importance weights assigned by the model. We then used KANALYZER to analyze the top 50 non-overlapping, non-redundant motifs, along with the full set of sequences and the reference sequence NC 045512.2. The results are summarized in Table 2. In cases where KANALYZER did not produce results for a specific motif, we assumed that it was located in a genomic region with high nucleotide variability (e.g., near residues 203-205 of the nucleocapsid protein [37]) or involved numerous successive deletions (e.g., the large 9-base SGF deletion in OR1ab [38]). To improve the signal for these motifs, we extended them to 30 nucleotides based on a consensus sub-sequence from the genomes where they were initially present. These extended motifs (ID 3, 8, 12, 18, and 38 in Table 2) were then analyzed using KANALYZER like the others. As shown on Table 2, the majority of the identified motifs were located in the coding regions of structural proteins, particularly the S protein. These motifs tended to involve missense mutations, which can have significant impacts on the infectivity, tropism, and pathogenesis of the virus even when few changes are involved [39]. Motif 1, located in the S glycoprotein, is an interesting example. It has a variation present in Beta variants and in 90% of Omicron variants that involves the K417N mutation. A second variation of motif 1, found in Gamma variants, involves the K417T mutation. Both mutations occur in the receptor binding domain (RBD) of S protein, which plays a crucial role in viral infection by interacting with the host ACE2 cell surface receptor. According to published reports, these mutations may potentially decrease binding ACE2 [40] and facilitate immune escape [41]. In contrast to the K417N/T mutations, the N501Y substitution found in the RBD-ACE2 interface was shown to result in one of the largest increases in ACE2 affinity conferred by a single RBD mutation [40]. This substitution, which is associated with the variation of motif 4, is present in several different variants, including Alpha, Beta, Gamma, and Omicron. According to Nelson et *al*. [42], the additional presence of the E484K mutation can further enhance virus binding to ACE2, while the presence of the K417N substitution can stabilize this binding. The combination of these mutations may result in the emergence of a mutant, whith the potential to evade host immune responses [42]. In addition, tests in individuals who received the Moderna or Pfizer-BioNTech SARS-CoV-2 vaccines suggest that the presence of the K417N, N501Y, and E484K mutations may result in a small but significant reduction in viral neutralization, potentially impacting the effectiveness of these vaccines against certain variants [43]. KEVOLVE highlighted several other notable mutations in the S protein, including the P681H and P681R substitutions. P681H is present in the sequences of both Alpha and Omicron variants, and its proximity to the furin protease cleavage site is thought to increase the cleavage of the S protein, potentially contributing to the rapid transmission of these variants [44]. This mutation was suggested to enhance SARS-CoV-2 infectivity [45]. The P681R substitution, which is highly conserved in the Delta and Kappa variants, appears to be associated with enhanced fusogenicity and pathogenicity [8]. The Omicron variant is distinguished by the N679K substitution, which is associated with motif 28 and also located near the furin cleavage site [46]. When combined with P681H, both substitutions allow for the inclusion of basic amino acids near the furin cleavage site, facilitating the partition of the S protein into S1 and S2 subunits and enhancing virus fusion and infection [47]. Among other notable mutations in the S protein, KEVOLVE identified the double Del156-157 and R158G substitution (highlighted by motif 3), which are located in the N-terminal domain (NTD) of the protein and are unique to the Delta variant. These mutations, known as vaccine breakthrough mutations [48], may potentially contribute to enhanced transmissibility or reduced sensitivity to pre-existing neutralizing antibodies [49]. Motif 3 also allowed the identification of the W152C and E154K mutations, which are present in more than 98% of Epsilon variants and ≈ 80% of Kappa variants. The W152C mutation, in particular, is correlated with the S13I mutation associated with motif 43, which together have important biological consequences that may allow immune evasion [50]. According to [50], mass spectrometry and structural studies showed that the S13I and W152C mutations resulted in a complete loss of neutralization for 10 of 10 NTD-specific monoclonal antibodies, due to the remodeling of the NTD antigenic supersite by the shift of the signal peptide cleavage site and the formation of a new disulfide bond.

**Table 2.**
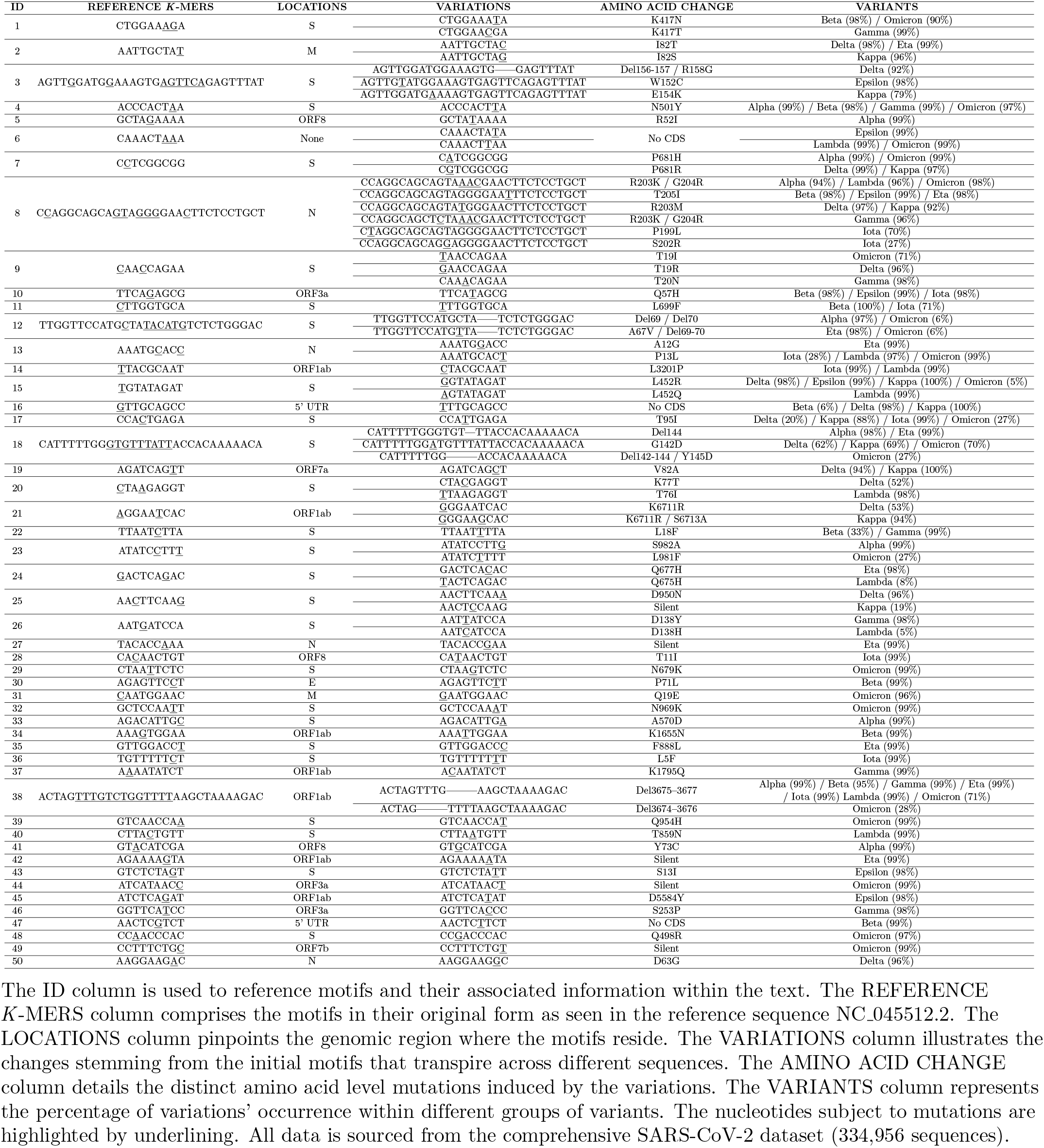
Mutational landscape of the motifs identified by KEVOLVE.

Other examples of mutations that affect the ability of SARS-CoV-2 to bind to specific antibody molecules (antigenicity) include the L18F, T19R/I, and T20N substitutions, which are highlighted by motifs 9 and 22. L18F is found in the Gamma variant and in ≈ 35% of Beta genomes. T19R and T19I are present in 96% of Delta variants and 71% of Omicron variants, respectively, while T20N is a Gamma-specific mutation. Epitope binding of 41 NTD-specific monoclonal neutralizing antibodies (mAbs) identified six antigenic sites, one of which, termed the “NTD supersite”, is recognized by all known NTD-specific mAbs and consists of residues 14-20, 140-158, and 245-264 [51]. The mutations associated with motifs 9 and 22 therefore include substitutions close to these antigenic regions of the NTD, including L18F, which is known to reduce neutralization by some antibodies [52]. A last example of motif located in the S protein identified by KEVOLVE that involves major impacts on the characteristics of SARS-CoV-2 is motif 14. A first variation of this motif, present in Delta, Epsilon, Kappa, and a minority of Omicron variants (5%), involves the L452R substitution. Located in the spike RBD which interacts directly with ACE2, this mutation was shown to increase spike stability, viral infectivity, viral fusogenicity, and viral replication [53]. The L452Q substitution, which is present in the Lambda variant, appears to be correlated with the T76I mutation associated with motif 20. These specific mutations are major contributors to the increased infectivity of the Lambda variant compared to other variants [54].

Regarding the mutations of interest associated with the motifs identified by KEVOLVE outside the S protein are I82T and I82S which are located in the M protein. M protein is highly conserved with low mutation rates and is a key element in virion morphogenesis and assembly, facilitating the release of viral particles from host cells and enhancing glucose transport during replication [55]. The I82T mutation, found in Delta and Eta variants, was suggested to enhance viral replicative fitness by altering cellular glucose uptake [56]. The I82S mutation, which is currently unique to Kappa, has not yet been well studied for its effects on SARS-CoV-2 [57]. Motif 8, located in the highly immunogenic and abundantly expressed N protein, is a last relevant example of a motif associated with mutations of interest. KANALYZER’s analysis of this motif has identified variations in that region that involve P199L, S202R, R203K/M, G204R, and T205I, at least one of which is found in every major natural variant [58]. The R203K/G204R mutation, which is present in the majority of Alpha, Gamma, Lambda, and Omicron variants, was shown to confer replication advantages likely related to ribonucleocapsid (RNP) assembly, and to be associated with increased infectivity, adaptability, and virulence of SARS-CoV-2 [59]. The R203M mutation, present in Delta and Kappa, as well as the S202R mutation present in ≈ 27% of Iota variants, were shown to increase viral infectivity by ≈ 50-fold [58]. Addition of the P199L mutation (present in ≈ 70% of Iota variants) to S202R and R203K/M increases transmissibility by four to seven times and enhances luciferase activity, which is positively correlated with the more efficient assembly of virus-like particles and more effective mRNA delivery [58]. Overall, the highly variable region of residues 203-205 in the N protein of SARS-CoV-2, which includes the T205I substitution specific to Beta, Epsilon, and Eta, was associated with increased replication and pathogenicity [37]. The motif analysis reports generated by KANALYZER and the accession numbers of the sequences used in our study are available on our GitHub directory (https://github.com/bioinfoUQAM/KEVOLVE). In addition, all identified mutations were manually confirmed using resources found at https://covdb.stanford.edu/variants/ and https://covariants.org/.

### Motifs identified by KEVOLVE/KANALYZER as genomic signature of SARS-CoV-2 variants

In the comparative study, we used KEVOLVE to identify motifs that discriminate between different classes of SARS-CoV-2 variants. We then selected the top 50 non-overlapping and non-redundant motifs determined by the importance weights assigned by the model. These 50 motifs were subsequently input into KANALYZER to characterize and identify their variations within the different SARS-CoV-2 variant groups (Column “VARIATIONS” of Table 2).” In total, we obtained 125 motifs and their associated variations, which are represented in the form of a cluster map (Fig 3). This map illustrates the frequency of absence/presence of each motif across different SARS-CoV-2 variants. Although these motifs were identified by KEVOLVE from a training subset of 2,500 sequences, the frequencies shown in Fig 3 are computed from the entire dataset of 334,956 sequences. By examining the columns, it is possible to identify different profiles and clusters of absence/presence of motifs specific to various variants. For example, Omicron has a cluster of 7 motifs that are unique to this variant (located in the lower left of the cluster map), with the exception of the ATCATAACT motif, which is also present in Iota. Towards the middle of the cluster map, we can see a second cluster of 7 motifs that appear in all variants except Omicron. These two Omicron-specific clusters contribute to its distance from the other SARS-CoV-2 variants.

**Fig 3.**
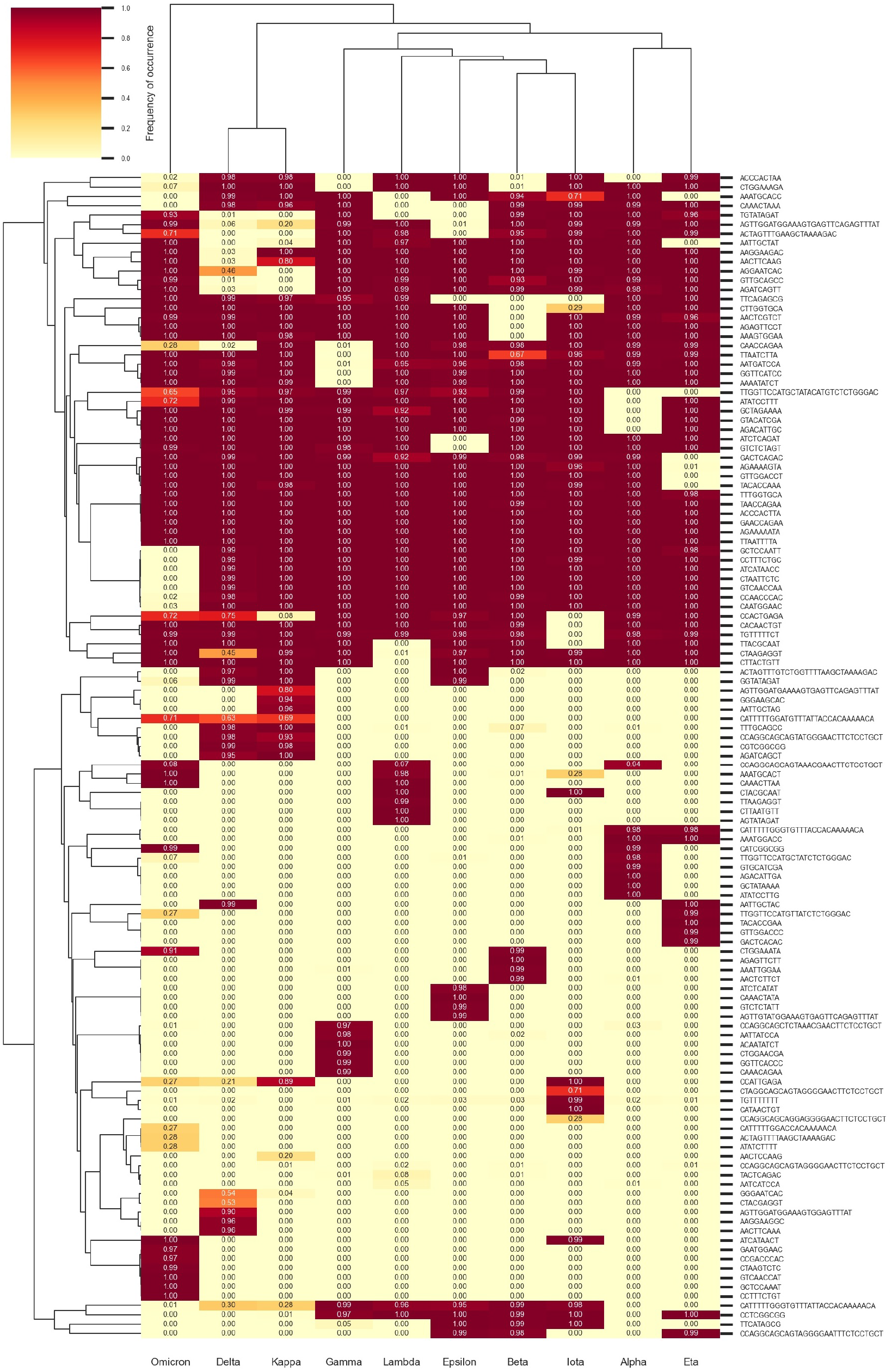
Cluster map of motif occurrence frequency according to SARS-CoV-2 variants.

In summary, this figure illustrates KEVOLVE’s ability to identify motifs in temporally conserved regions starting with a limited set of sequences and to generalize to a larger dataset of sequences collected since the start of the COVID-19 pandemic. The identified motifs provide genomic signatures that can be used to generate peptide or oligonucleotide libraries for rapid and accurate detection of listed pathogens with tools such as VirScan [60] or to design specific primer sets for the classification of SARS-CoV-2 variants with artificial intelligence [61]. These approaches, which use models built from a restricted number of motifs and sequences, can efficiently classify large sets of sequences, which is crucial during major viral outbreaks where swift identification of the virus’ taxonomic classification and genomic sequence origin is necessary for effective strategic planning, containment, and treatment [10]. In addition, the identified genomic signatures, along with the reports generated by KANALYZER, provide valuable insights that can help understand the viral evolution and transmission, the mechanisms through which the virus causes disease, and the development of treatments and vaccines. These approaches, which use models built from a restricted number of motifs and sequences, can efficiently classify large sets of sequences, which is crucial during major viral outbreaks where swift identification of the virus’ taxonomic classification and genomic sequence origin is necessary for effective strategic planning, containment, and treatment [10].

Finally, although our current approach operates within a closed classification framework, which is limited to the classes defined by the training sequence dataset, we plan to extend it to an open classification context. To achieve this, we propose a strategy to calculate the distance between each new sequence and the existing genomic signature profiles, generated in the cluster map (Fig 3). By using an appropriate distance threshold, we can identify sequences that are significantly distant from known signatures, potentially indicating a new variant. Thresholds can be determined by leveraging the knowledge of distances between genomic signature profiles of different known variants. This method could be based on distance metrics such as Euclidean distance, Manhattan distance, or even a normalized distance based on *k*-mer similarity. Furthermore, to make our approach more flexible and adaptable to new variants, we could also implement an incremental learning mechanism. In this way, each time a new variant is identified above a certain support threshold, the associated sequences could be integrated into the initial training set, and the model would be retrained to account for this new information. This would allow our model to learn and progressively adjust its parameters based on the newly encountered sequences. This approach could facilitate the detection of new variants and enable regular model updates with the integration of new sequences associated with emerging variants.

## Conclusion

In this study, we compared the performance of the machine learning-based tools KEVOLVE and CASTOR-KRFE with statistical tools specialized in identifying discriminative motifs in unaligned sequence sets for the classification of SARS-CoV-2 variants. Overall, the models based on the motifs identified by KEVOLVE outperformed the models based on the motifs identified by the statistical tools, while using a lower number of motifs. Models based on STREME motifs achieved the second-best performance (slightly below KEVOLVE), but these models require the use of twice as many motifs. The drop in performance was mainly due to prediction errors for Beta, Kappa, and Lambda variants. CASTOR-KRFE obtained the third-best performance with models based on 10 times fewer motifs than STREME, as the tool only identifies a single subset of motifs, unlike the others. The prediction errors of the CASTOR-KRFE models are associated with the same variants as those of STREME, but they are more pronounced. Finally, the weakest performances were associated with the MEME OOPS/ZOOPS models, with many more errors for the same variants than STREME and CASTOR-KRFE. This study also demonstrated that KEVOLVE and CASTOR-KRFE are able to handle multi-class sets, rather than being limited to binary sets like some other tools. This is an important advantage when analyzing organisms such as SARS-CoV-2, which are constituted of multiple classes of viral variants.

Subsequently, we analyzed the motifs identified by KEVOLVE using KANALYZER, a new extension based on pairwise alignment and parallel computing. This analysis allowed us to identify variations of the discriminative motifs in different classes of SARS-CoV-2 variants, including their frequency, genomic localization, and mutation at the amino acid level. This analysis, performed on all 334,956 sequences belonging to the 10 major variant classes defined by the WHO, showed that the majority of the motifs identified by KEVOLVE were located in structural proteins, with a particular focus on the S protein. The motifs and variations identified were linked to known mutations previously reported in the literature, which are assumed to affect key characteristics of the virus such as infectivity, pathogenicity, tropism, transmission, and evolution. In conclusion, this study demonstrates the utility of KEVOLVE as a robust tool for identifying discriminative motifs of SARS-CoV-2 variants. These motifs provide genomic signatures that can be used to construct oligonucleotide libraries or to build artificial intelligence models for rapid and accurate pathogen detection. Furthermore, KANALYZER allows the analysis of motifs identified by KEVOLVE, providing valuable insights into the biological properties of viruses and viral gene products that serve as targets for the development of vaccines or antiviral therapy.

